# Circadian and diel regulation of photosynthesis in the bryophyte *Marchantia polymorpha*

**DOI:** 10.1101/2022.01.11.475783

**Authors:** David Cuitun-Coronado, Hannah Rees, Joshua Colmer, Anthony Hall, Luíza Lane de Barros Dantas, Antony N. Dodd

## Abstract

Circadian rhythms are 24-hour biological cycles that align metabolism, physiology and development with daily environmental fluctuations. Photosynthetic processes are governed by the circadian clock in both flowering plants and some cyanobacteria, but it is unclear how extensively this is conserved throughout the green lineage. We investigated the contribution of circadian regulation to aspects of photosynthesis in *Marchantia polymorpha,* a liverwort that diverged from flowering plants early in the evolution of land plants. First, we identified in *M. polymorpha* the circadian regulation of photosynthetic biochemistry, measured using two approaches (delayed fluorescence, PAM fluorescence). Second, we identified that light-dark cycles synchronize the phase of 24 h cycles of photosynthesis in *M. polymorpha*, whereas the phases of different thalli desynchronize under free running conditions. This might also be due to masking of the underlying circadian rhythms of photosynthesis by light-dark cycles. Finally, we used a pharmacological approach to identify that chloroplast translation might be necessary for clock control of light harvesting in *M. polymorpha.* We infer that the circadian regulation of photosynthesis are well-conserved amongst terrestrial plants.

## Introduction

The rotation of the Earth on its axis causes 24-hour cycles in environmental conditions such as light and temperature. These diel cycles are thought to have selected for the evolution of circadian clocks within several Kingdoms of life (Dodd *et al.* 2005, Eelderink-Chen *et al.* 2021, Millar 2016, Ouyang *et al.* 1998, Spoelstra *et al.* 2016). Circadian rhythms are self-sustaining biological cycles that have a period of about 24 h, which are thought to provide a biological measure of the time of day. These rhythms coordinate and sequence processes around the diel cycle, whilst ensuring that responses to environmental cues are appropriate for the time of day.

Circadian rhythms in plants are generated by a molecular oscillator that is formed from a series of transcription-translation feedback loops, which has a predominance of negative feedback steps (Hsu and Harmer 2014). The oscillator is entrained to environmental cues such as light and temperature, which align the phase of the rhythm with the day/night cycle so that it provides an accurate biological measure of the time of day. There is circadian regulation of a variety of key aspects of the physiology of flowering plants, such as stomatal opening, light harvesting, CO_2_ fixation and starch metabolism (Dakhiya and Green 2022, Dakhiya *et al.* 2017, Dodd *et al.* 2005, Graf *et al.* 2010). Furthermore, the circadian oscillator contributes to the photoperiodic control of flowering time (Hayama and Coupland 2003, Park *et al.* 1999) and the fitness of plants (Dodd *et al.* 2005, Green *et al.* 2002). This means that circadian regulation has a pervasive influence upon physiology, metabolism, and development (Hotta *et al.* 2007, Millar 2016), reflecting the close functional relationship between plants and their fluctuating environments.

Circadian regulation of photosynthesis has been reported in a variety of flowering plants, including *Arabidopsis thaliana, Hordeum vulgare, Zea mays, Phaseolus vulgaris, Kalanchoë daigremontiana* and *Mesembryanthemum crystallinum* (Dakhiya *et al.* 2017, Davies and Griffiths 2012, Dodd *et al.* 2003, Dodd *et al.* 2004, Dodd *et al.* 2005, Gould *et al.* 2009, Hennessey and Field 1991, Wyka *et al.* 2005). Aspects of photosynthetic physiology that are circadian regulated include the rates of CO2 assimilation and oxygen evolution, and photosynthetic light harvesting reported using chlorophyll fluorescence analysis (Dakhiya *et al.* 2017, Hennessey and Field 1991, Litthauer *et al.* 2015, Schweiger *et al.* 1964). There are also circadian rhythms in stomatal opening, which are functionally separable from rhythms of CO_2_ assimilation, because rhythms of assimilation continue when the intercellular partial pressure of CO_2_ is held at a constant level to remove the influence of rhythmic stomatal opening upon assimilation (Hennessey and Field 1991). In addition to flowering plants, aspects of metabolism associated with photosynthesis are circadian regulated in some cyanobacteria and algae (Cano-Ramirez *et al.* 2018, Mitsui *et al.* 1986, Samuelsson *et al.* 1983, Schneegurt *et al.* 1994, Schweiger *et al.* 1964). Despite this evidence for the circadian regulation of photosynthesis, the mechanisms and evolution of this process remain less well understood.

The pervasiveness of circadian regulation of photosynthesis within flowering plants and unicellular photosynthetic organisms might suggest that this process is conserved throughout multicellular plant life, with a potentially ancient evolutionary origin. One approach to examine this idea is to investigate plant species that diverged from flowering plants at a relatively early stage of land plant evolution, because this can establish whether the mechanism might originate from a common ancestor. As a model to investigate this question, we selected the bryophyte liverwort *Marchantia polymorpha*. *M. polymorpha* is thought to have diverged from flowering plants about 400 million years ago (Delwiche and Cooper 2015, Kohchi *et al.* 2021), with liverwort-like plants occurring earlier in the fossil record than other land plant forms such as vascular plants (Edwards *et al.* 1995). *M. polymorpha* is a useful model to investigate questions concerning the evolution of circadian regulation because it has a circadian oscillator that shares some components with flowering plants (Lagercrantz *et al.* 2021, Linde *et al.* 2017), there is circadian regulation of a subset of the transcriptome and the position of its thallus lobes (Lagercrantz *et al.* 2021, Lagercrantz *et al.* 2020), and it can be used in experimental designs comparable to models such as Arabidopsis. Whilst circadian rhythms and some aspects of circadian clock structure have been identified in *M. polymorpha*, it is not yet known whether the regulation of photosynthesis represents a conserved output from its circadian system.

Here, we used *M. polymorpha* as a tool to investigate three questions concerning the conservation of circadian regulation of photosynthesis in multicellular plants. First, we investigated whether there is circadian regulation of photosynthesis in *M. polymorpha,* to determine whether this process is conserved in a species that diverged from vascular plants at a relatively early stage of land plant evolution. Second, we investigated whether a circadian clock in *M. polymorpha* might influence the daily regulation of photosynthesis under cycles of light and dark, to determine whether circadian regulation might contribute to the timing of photosynthesis under diel cycles. Finally, we investigated whether pharmacological approaches might provide insights into the mechanisms that underlie the circadian rhythms of photosynthesis.

## Results

### Circadian rhythms of photosynthesis in M. polymorpha

We used two proxies of photosynthetic activity to investigate circadian rhythms of photosynthesis in *M. polymorpha.* These were delayed chlorophyll fluorescence (DF), and prompt fluorescence (PF) measured using the pulse amplitude modulation (PAM) method (Dakhiya and Green 2022, Gould *et al.* 2009, Maxwell and Johnson 2000). When illuminated leaves are transferred to darkness, the potential for electrons to move down the electron transport chain is removed, and energy within the system is instead re-emitted as light and heat. The light emitted within the nanosecond range after the lights turn off is referred to as prompt fluorescence (Dakhiya and Green 2022, Maxwell and Johnson 2000). Light emitted within the millisecond to second range following lights-off is referred to as DF (Gould *et al.* 2009, Rees *et al.* 2019, Strehler and Arnold 1951). Both DF and certain features of PF are circadian-regulated in a range of flowering plants and some algae (Cano-Ramirez *et al.* 2018, Dakhiya and Green 2022, Gould *et al.* 2009, Gyllenstrand *et al.* 2014). One exception appears to be spruce trees *(Picea abies),* where the DF rhythm damps rapidly in accordance with a rapidly damping circadian oscillator (Gyllenstrand *et al.* 2014). Features of chlorophyll fluorescence that oscillate with a circadian rhythm include the apparent quantum yield of PSII (YII), a measure of non-photochemical quenching of chlorophyll fluorescence within PSII (NPQ), and the estimated photosynthetic electron transport rate (ETR) (Cano-Ramirez *et al.* 2018, Dakhiya and Green 2022, Dakhiya *et al.* 2017). Taken together, rhythms of DF and PF describe circadian regulation of processes of photochemistry over time, as greater energy loss as heat (NPQ) and light (DF) indicate lower quantum yields (Murchie and Lawson 2013).

We investigated whether free-running circadian rhythms of DF occur in *M. polymorpha.* For these experiments, plants were investigated under conditions of either continuous light, which was interrupted hourly by darkness for the measurement of DF, or under conditions of continuous darkness, interrupted hourly by a 5 minute light pulse to allow subsequent DF measurement. We named these conditions free running light conditions (FRL) and free running dark conditions (FRD), respectively. We also investigated DF responses under cycles of 12 h light and 12 h darkness, with the light and dark periods interrupted at hourly intervals, for 5 minutes, for the measurement of DF. We named these conditions zeitgeber cycles (ZTC). Rhythmic thalli were identified using Metacycle (Wu *et al.* 2016) (*q* < 0.001; period between 18 and 34 hours). Under FRL conditions, 63% of thalli had circadian rhythms of DF that persisted for 4-5 days, and damped over time (Fig 1a, b; Table 1; Fig. S1; Metacycle *q* < 0.001). The mean period of these rhythms under FRL conditions was 25.50 h ± 0.52 h (mean ± s.e.m.). Under FRD conditions, 78% of thalli had circadian rhythms of DF (Table 1; Metacycle *q* < 0.001). However, there was considerable variability between the rhythms of individual thalli, so the mean DF signal from all rhythmic thalli had little overt rhythmicity (Fig. 1c, d; Fig. S2). In contrast, under zeitgeber cycles, 100% of thalli were rhythmic (Metacycle *q* < 0.001), and the rhythms had better phase synchronization between individual thalli (Fig. 1e, f; Fig. S3).

**Figure 1.**
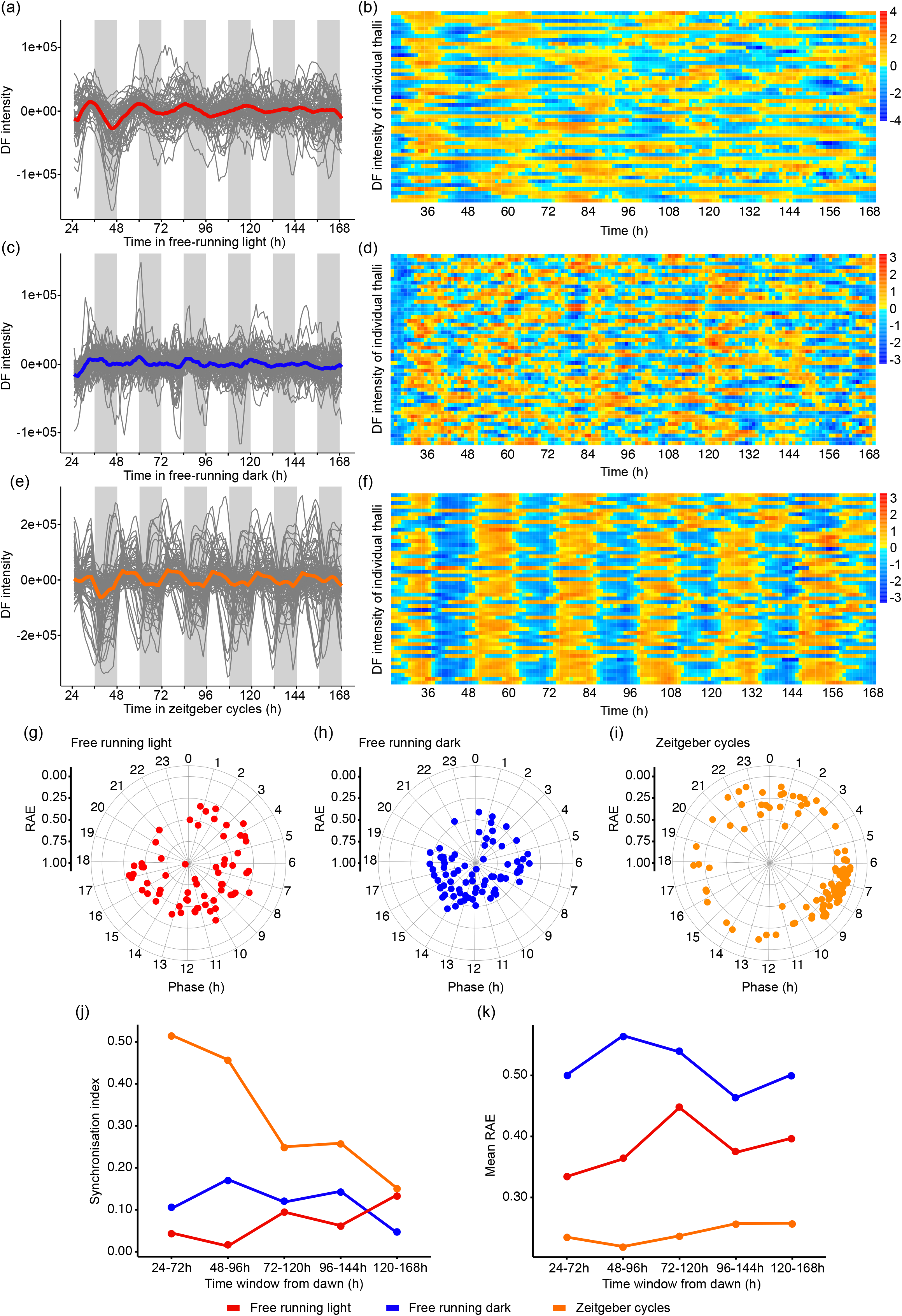
Circadian and diel rhythms of delayed fluorescence of *Marchantia polymorpha* thalli under (a, b) free running light, (c, d) free running dark, and (e, f) zeitgeber cycle conditions. (a, c, e) indicate DF signal from thalli categorized as rhythmic after filtration with Metacycle (*q*-value < 0.001; period between 18 h and 34 h). The mean is the thick coloured line and individual replicate thalli are fine grey lines. (b, d, f) Heat maps summarize DF intensity of 50 individual and randomly selected thalli across two experimental repeat experiments under (b) free running light, (d) free running dark and (f) zeitgeber cycle conditions. Data in (a, c, e) are smoothed with a 5-hour moving average, baseline and amplitude de-trended, and normalized to the mean signal. On heat maps, colour values are scaled per row using Z-score normalization. (g-i) Phase of replicate thalli relative to dawn (0), calculated using fast Fourier transform-non linear least squares method (FFT-NLLS). Radial scale shows relative amplitude error (RAE) of each thallus, with RAE=0 at the exterior and RAE=1 at the centre of the circles, respectively. (j) Comparison of relative synchronization between replicate thalli under each light condition, using the order parameter within the Kuramoto phase oscillator model to provide a synchronization index. Greater values indicate better phase synchrony between replicates. (k) Comparison of RAE change over time under each light condition. Data derive from 100 individual thalli, across two independent experimental replicates, for each condition.

**Table 1.**
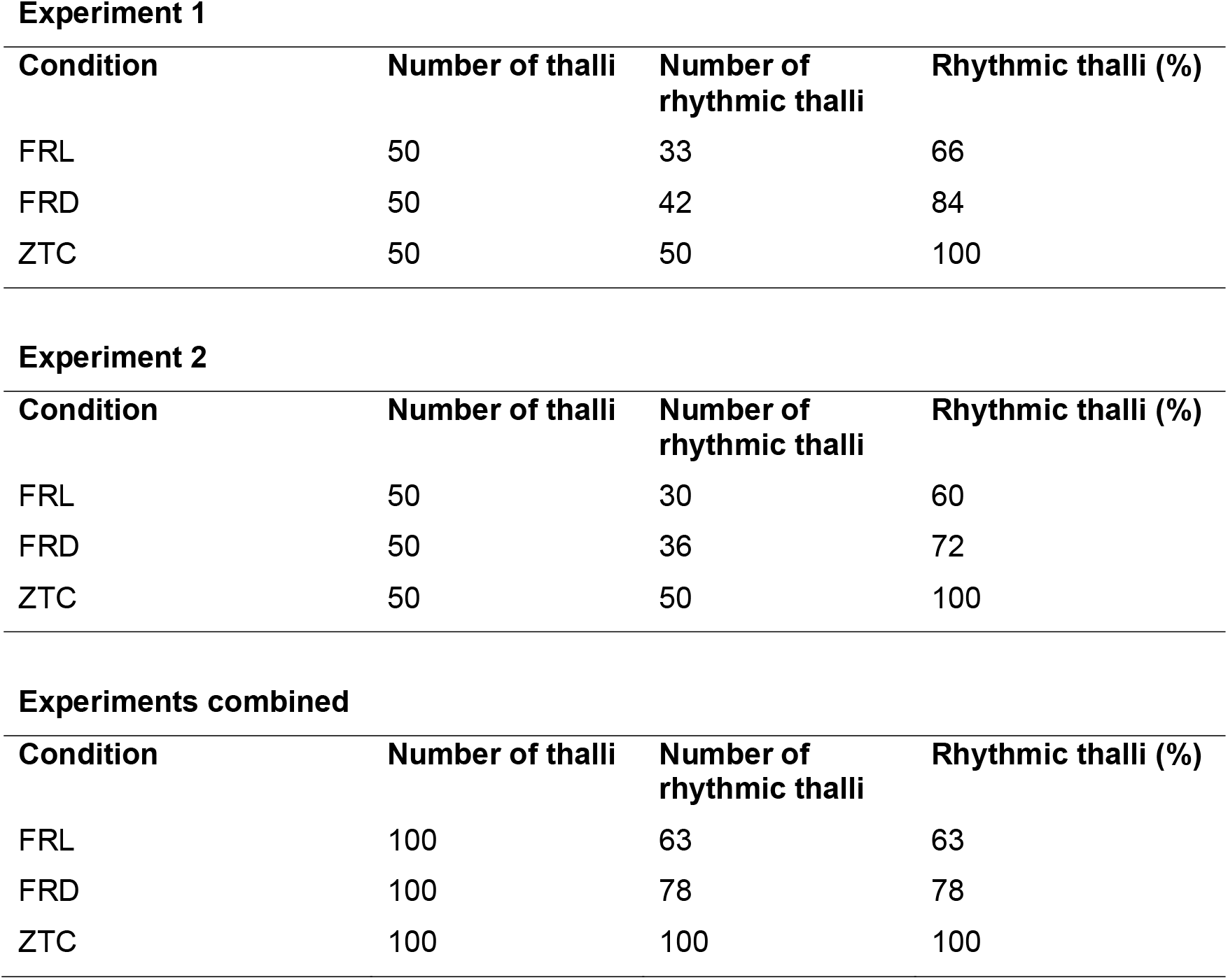
Comparison of number of *M. polymorpha* thalli having a circadian or diel rhythm of delayed fluorescence under each light condition, from two independent experiments. Data were classified as rhythmic when the Metacycle q-value was < 0.001 and period lengths between 18 h and 34 h. The abbreviations FRL, FRD and ZTC represent free running conditions of light, free running conditions of darkness, and zeitgeber cycles (light/dark cycles), respectively.

**Experiment 1**
| Condition | Number of thalli | Number of rhythmic thalli | Rhythmic thalli (%) |
| --- | --- | --- | --- |
| FRL | 50 | 33 | 66 |
| FRD | 50 | 42 | 84 |
| ZTC | 50 | 50 | 100 |

**Experiment 2**
| Condition | Number of thalli | Number of rhythmic thalli | Rhythmic thalli (%) |
| --- | --- | --- | --- |
| FRL | 50 | 30 | 60 |
| FRD | 50 | 36 | 72 |
| ZTC | 50 | 50 | 100 |

**Table 1.** Comparison of number of *M.
| Condition | Number of thalli | Number of rhythmic thalli | Rhythmic thalli (%) |
| --- | --- | --- | --- |
| FRL | 100 | 63 | 63 |
| FRD | 100 | 78 | 78 |
| ZTC | 100 | 100 | 100 |

Rapid damping of a rhythm over time (Fig. 1a, red line) can suggest the desynchronization of rhythms of individual replicates (Paajanen *et al.* 2021). We hypothesized that such desynchronization might occur for DF in *M. polymorpha* under our conditions. This is supported by the considerable range of phases across replicate thalli under both FRL and FRD conditions (Fig. 1g, h), which contrasts zeitgeber cycle conditions where the phase of many replicates clustered to the end of the subjective day (6 – 10 h; Fig. 1i). To investigate this further, we calculated the synchrony of phase between individual replicates, using the order parameter *r* within the Kuramoto phase oscillator model (Kuramoto 1984, Muranaka and Oyama 2016). This identified greater phase synchrony under zeitgeber cycles compared with free running conditions (Fig. 1j), with phase synchrony weakening over time under zeitgeber cycles. We were interested in the level of phase synchrony of DF in *M. polymorpha* compared with other species, so also calculated also phase synchrony for DF in Arabidopsis, *Triticum aestivum* and *Brassica napus* from published data (Rees *et al.* 2019, Rees *et al.* 2021) (Dataset S1). This identified very high phase synchrony for replicates of Arabidopsis (*r* = 0.92) and *B. napus* (*r* = 0.96) under FRL conditions, compared with relatively low phase synchrony for both *T. aestivum* under FRL conditions (*r* = 0.33) and *B. napus* under FRD conditions (*r* = 0.47) (Dataset S1).

To investigate the robustness of the relatively desynchronized rhythms in *M. polymorpha* we used the parameter of relative amplitude error (RAE), which is derived from analysis of the data by fast Fourier transform-non linear least squares method (FFT-NLLS). RAE provides a measure of the robustness of the oscillation, representing the ratio of the amplitude error to the amplitude, with a value of 0 indicating a robust and non-damping rhythm, and a value of 1 indicating an absence of rhythmicity, or very noisy data. RAE of < 0.6 is sometimes considered indicative of rhythmicity (Alabadí *et al.* 2001, Alabadi *et al.* 2002, Para *et al.* 2007). FFT-NLLS analysis suggested relatively robust rhythms under zeitgeber cycles, and lower rhythmic robustness under FRL and FRD conditions (Fig. 1k). The mean RAE of rhythms of DF for thalli under FRL conditions was 0.46 ± 0.01, which suggests a lower robustness of the rhythm compared to rhythms of DF in *Arabidopsis* seedlings under the same conditions (RAE = 0.31 – 0.44, depending on accession) (Rees *et al.* 2021). Taken together, this suggests that whilst a proportion of thalli were rhythmic under FRL and FRD conditions (Table 1), the rhythm of DF varied considerably between thalli and had lower rhythmic robustness compared with under zeitgeber cycles (Fig. 1k). This also supports the notion that light input masks or maintains the daily rhythm of DF in *M. polymorpha.*

To extend our DF analysis, we also measured circadian rhythms of photosynthetic activity in *M. polymorpha* using the PAM chlorophyll fluorescence method. In this experiment, plants were maintained under conditions of constant blue actinic light, with alterations in these light conditions every two hours to measure fluorescence parameters (Cano-Ramirez *et al.* 2018). Under these conditions within the PAM instrument, there were circadian rhythms in the effective quantum yield of PSII (Y(II)) (Fig. 2a, b) and a measure of non-photochemical quenching of chlorophyll fluorescence within PSII (NPQ) (Fig. 2c, d). These rhythms had a period between 27 h and 28 h (28.10 h ± 0.15 h (Y(II)), 27.55 h ± 0.38 h (NPQ)) in *M. polymorpha* (Fig. 2b, d). These period estimates are longer compared with the period estimates obtained using DF (Fig. 1h). This might be caused by the different light conditions used with the two methods, because the period of some circadian-regulated processes in *M. polymorpha* depends upon the light intensity and spectrum (Linde *et al.* 2017). Alternatively, this might reflect differences in the preparation of thalli for the two methods, since thalli were grown directly from gemmae for DF imaging, whereas for PAM experiments the thalli were cut into squares to prevent hyponasty. Y(II) and NPQ oscillated with opposing phases (Fig. 2a, c), which is consistent with other reports (Cano-Ramirez *et al.* 2018), and expected because these measures represent competing fates for light energy absorbed by leaves.

**Figure 2.**
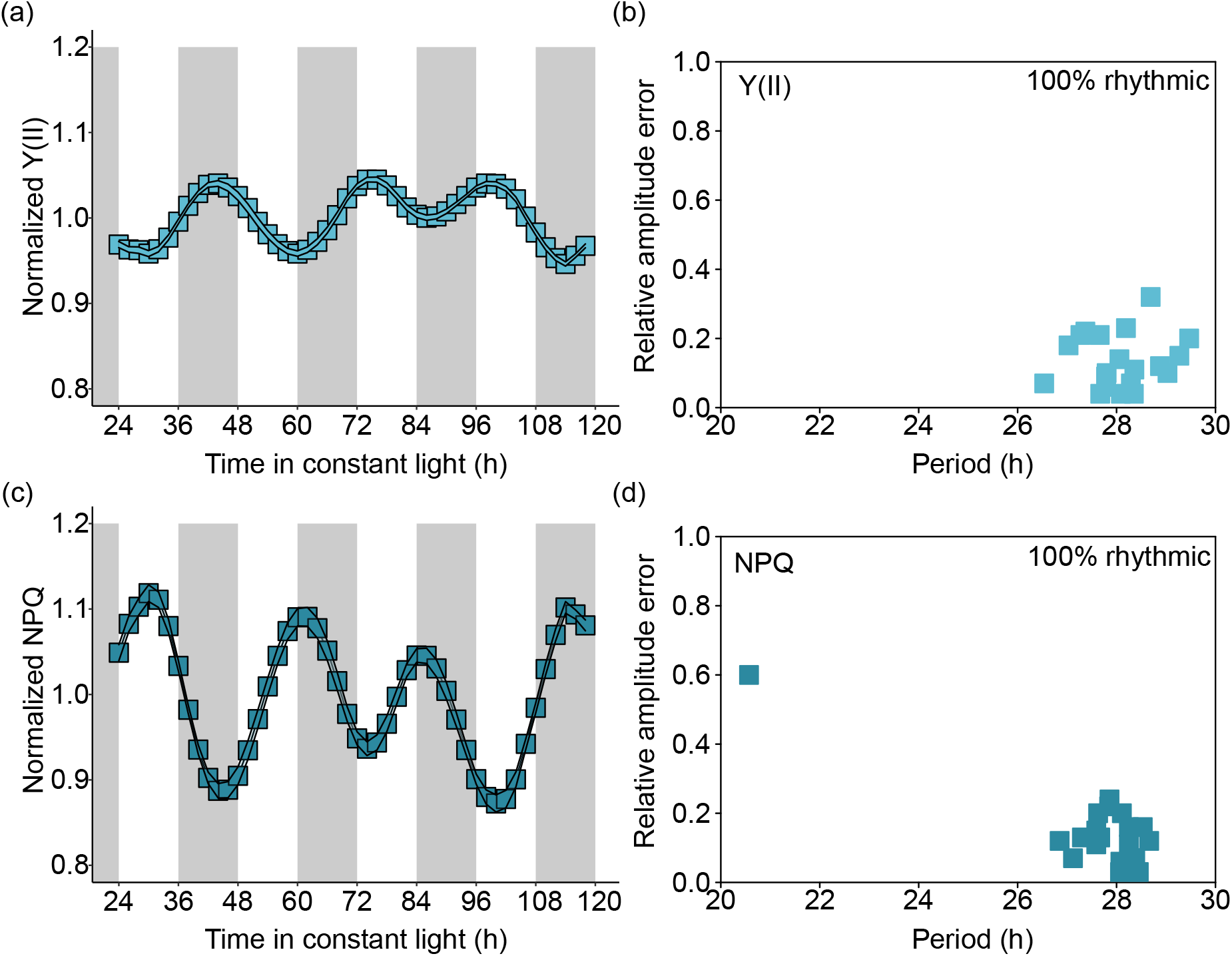
Circadian rhythms of modulated chlorophyll fluorescence in thalli of *Marchantia polymorpha.* Under free running conditions, time-course analysis of (a, b) the apparent quantum yield of PSII (Y(II)), and (c, d) non-photochemical quenching of PSII fluorescence (NPQ). (a, c) are the normalized signal level for each chlorophyll fluorescence parameter, processed using amplitude and baseline detrending and normalized to the mean (using Biodare2), and shaded grey areas indicate the subjective night. (b, d) compare the estimated period and relative amplitude error (RAE) of the rhythmic thalli, with period, RAE and the presence of rhythmicity quantified using FFT-NLLS (using Biodare2). n = 20 independent thalli.

Together with the measurements of DF, we conclude that free-running rhythms of several photosynthetic outputs occur in *M. polymorpha*. The DF data suggest that free-running rhythms are less robust and more variable under FRD conditions, but remain detectable in a proportion of thalli (Table 1). Under zeitgeber cycles, the robustness of individual rhythms (Fig. 1e, f, k) and relatively high phase synchrony between replicate thalli (Fig. 1j) suggests that under these conditions, the rhythm of DF is influenced strongly by the zeitgeber cycle. This indicates that under zeitgeber cycles, the underlying circadian oscillation (Fig. 1e) is either driven by the light condition, or the underlying circadian rhythm is masked by the light condition. This is consistent with the notion that the light/dark cycle has a strong effect upon the diel regulation of the *M. polymorpha* circadian oscillator and its physiological outputs (Linde *et al.* 2017).

### Contribution of circadian regulation in M. polymorpha to the timing of rhythms of delayed fluorescence under zeitgeber cycles

Because we identified that the zeitgeber cycle strongly influences the daily oscillation of DF in *M. polymorpha* (Fig. 1a-f), we wished to investigate the extent of clock control of the phase of delayed fluorescence oscillations in *M. polymorpha* under zeitgeber cycles. To achieve this, we used an experimental approach whereby the phase of the rhythmic process was monitored under a range of zeitgeber cycle lengths (known as T cycles; e.g. T24 = 12 h light / 12 h dark; T28 = 14 h light / 14 h dark, and so on). This can identify effects of circadian regulation upon the timing of a process under diel cycles, because the circadian oscillator can influence the phase of the process, producing systematic changes in the phase of either the entire waveform (Aschoff 1960, Eelderink-Chen *et al.* 2021, Roenneberg *et al.* 2005), or parts of the rhythm (Dodd *et al.* 2014). Using the zeitgeber cycles of DF described earlier, we identified oscillations of DF under diel cycles of T20 to T28 (Fig. 3a-d). The period length of DF matched the T cycle length closely (20.09 h ± 0.06 h (T=20), 22.44 h ± 0.07 h (T=22), 24.36 h ± 0.07 h (T=24) and 28.17 h ± 0.06 h (T=28), measured using FFT-NLLS in Biodare2). When the phase was normalized to the length of the T cycle, we found that the T cycle length had no significant influence upon phase between T20, T22 and T24, however, the phase was significantly later under the longest T cycle of T28 (Fig. 3e). In general, a circadian oscillator would be expected to cause relatively earlier phases under longer T cycles (Aschoff 1960), including in plants (Dodd *et al.* 2014). This did not occur within our experiment (Fig. 3e), where instead the phase followed the zeitgeber cycle length (Fig. 3a-e). The dependency of the period and phase of DF upon the T cycle length suggests that any underlying circadian clock control was masked by the zeitgeber cycle. In circadian biology, masking is the process whereby the apparent coupling of an observable rhythm to a zeitgeber leads to a shared period and phase between the observed rhythm and the zeitgeber, independent of any clock control (Aschoff 1999, Roenneberg *et al.* 2005). Therefore, it appears that the interaction between the cycle of light and dark and the biochemical processes of light harvesting tend to conceal the underlying effect of circadian regulation under these experimental conditions.

**Figure 3.**
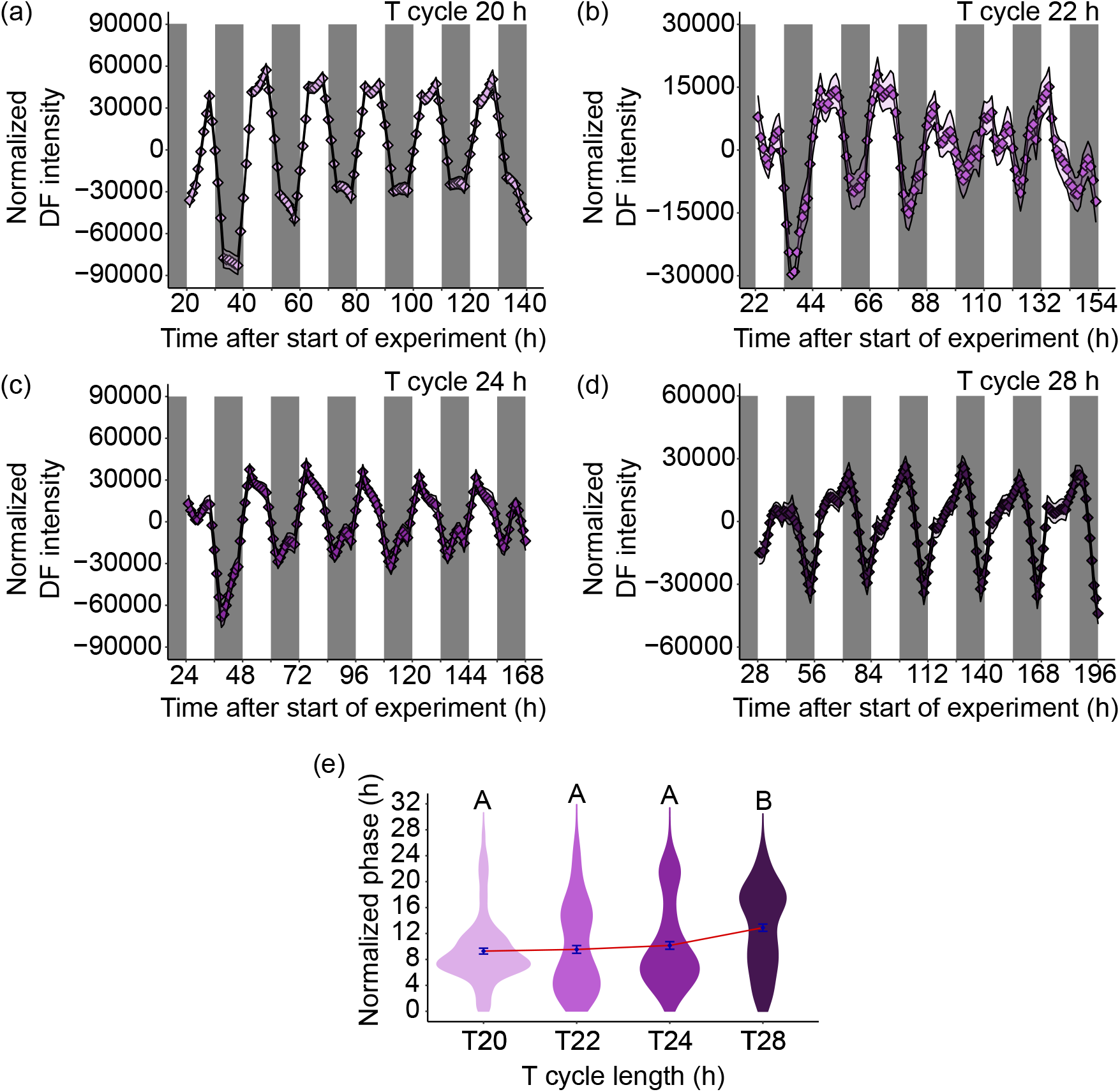
Analysis of the contribution of circadian regulation to the rhythms of delayed chlorophyll fluorescence under zeitgeber cycles. DF rhythms were quantified under T cycles of lengths of (a) 20 h, (b) 22 h, (c) 24 h and (d) 28 h, with the mean intensity plotted. (e) The phase was estimated from each time-series and normalized to a 24 h T cycle length, to evaluate the phase relationship between the rhythm of DF and the zeitgeber cycle. A strong contribution of the circadian clock to the timing of the rhythm under zeitgeber cycles would be expected to drive a relatively earlier phase under longer T-cycles, which (e) is not the case. (a-d) Shaded areas indicate dark periods. n = 150 thalli per T cycle (T20: 126 of 150 points rhythmic; T22: 127 of 150 points rhythmic; T24: 122 of 150 points rhythmic T28: 129 of 150 points rhythmic). Phase estimates obtained using FFT-NLLS, with data amplitude and baseline detrended. Only rhythmic thalli were included in FFT-NLLS analysis. In (e), different letters indicate significantly different mean phases (one-way ANOVA, where p < 0.05 based on Tukey’s post-hoc test).

### Abrogation of circadian rhythms of photosynthesis in M. polymorpha by a potential inhibitor of chloroplast translation

Whilst the canonical circadian oscillator underlies the daily timing of photosynthesis (Dodd *et al.* 2004, Dodd *et al.* 2005), the mechanisms by which the circadian oscillator regulates photosynthesis remain poorly understood. One approach to identify potential cellular processes associated with circadian rhythms of photosynthesis is to use a pharmacological strategy, because several chemicals are known to inhibit specific processes within chloroplasts. We combined a pharmacological approach with the monitoring of circadian rhythms of chlorophyll fluorescence to identify candidate processes associated with the circadian regulation of photosynthesis.

First, we tested whether an inhibitor of chloroplast translation can attenuate circadian rhythms of chlorophyll fluorescence in *M. polymorpha.* Lincomycin is an antibiotic that binds to the 50S subunit of bacteria-like ribosomes (Chang and Weisblum 1967) and is thought to inhibit chloroplast but not mitochondrial protein synthesis (Sullivan and Gray 1999, Zhao *et al.* 2018). We supplemented the growth media with a range of lincomycin concentrations (25 μg ml^-1^, 50 μg ml^-1^ and 100 μg ml^-1^, and a vehicle control), and monitored rhythms of photosynthesis using the PAM fluorescence method (Fig. 4a-l; Fig. S4a-d). There was a dose-dependent effect of lincomycin on the amplitude of rhythms of Y(II) and NPQ under free running light conditions (Fig. 4f, l; Fig. S4b, d), such that the amplitudes of these parameters were reduced by greater lincomycin concentrations (Fig. 4f, l; Fig. S4b, d)). Although the change was statistically significant, there was some variation between the experimental repeats (Fig. 4f, l; Fig. S4b, d). Furthermore, the RAE of circadian rhythms of Y(II) was significantly greater in the presence of lincomycin compared with the vehicle control (Fig. 4e; Fig. S4a). The effect of lincomycin on the RAE of rhythms of NPQ was less consistent across two experiments (Fig. 4k; Fig. S4c), suggesting that lincomycin might have a greater effect upon the rhythmic robustness of Y(II) compared with NPQ. These measures of photosynthesis are ratiometric, so the decreased amplitude is unlikely to be caused by a lower fluorescence signal in the presence of lincomycin. Whilst we cannot exclude the possibility that lincomycin in *M. polymorpha* affects processes other than plastid translation, our data suggest that chloroplast translation might be required to maintain circadian rhythms of photosynthetic light harvesting.

**Figure 4.**
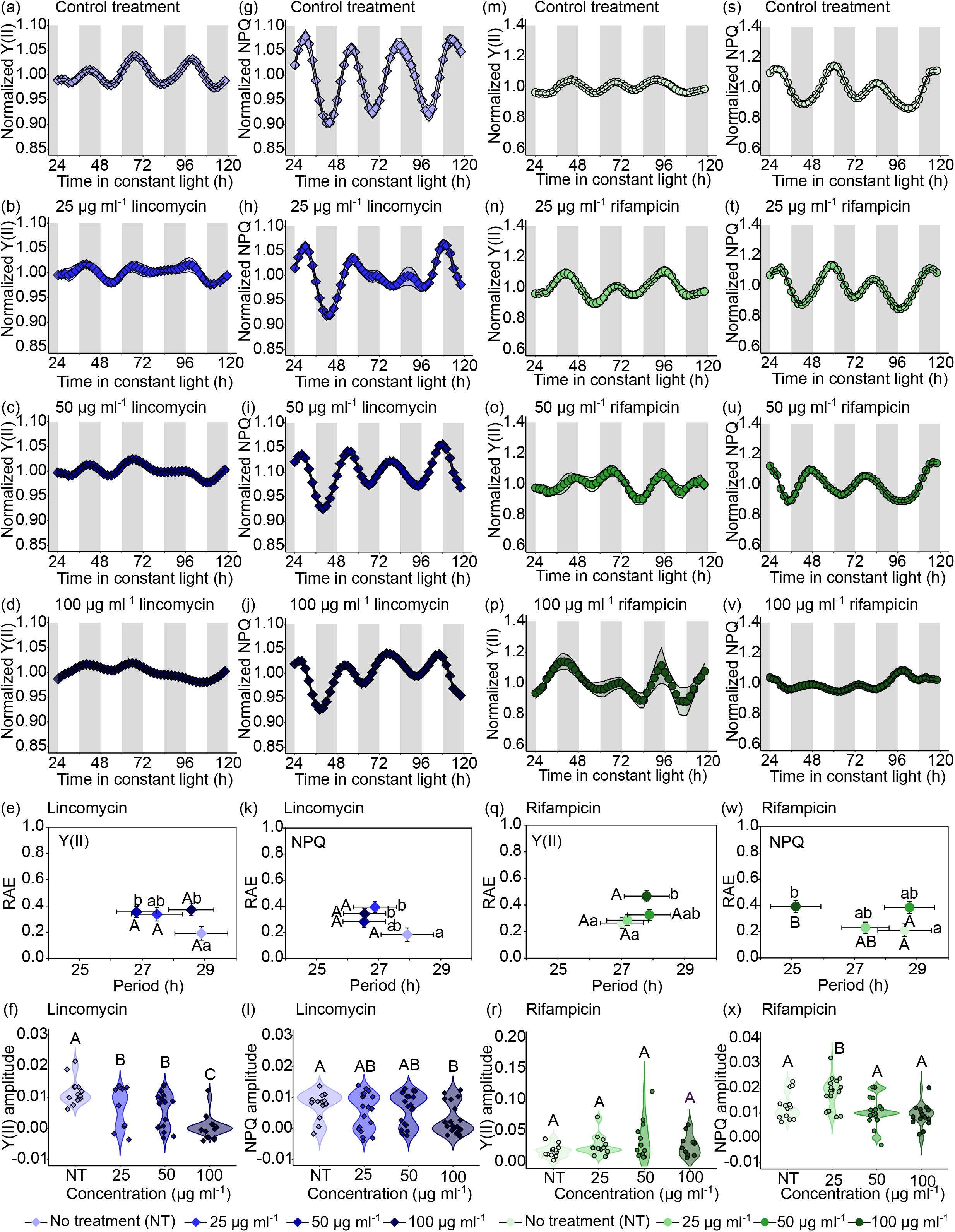
Amplitude of circadian rhythm of chlorophyll fluorescence parameters is reduced by the inhibitor of chloroplast translation lincomycin. (a-f) Effect of range of concentrations of lincomycin upon the circadian oscillation of the apparent quantum yield of PSII (Y(II)). (g-l) Effect of range of concentrations of lincomycin upon the circadian oscillation of non-photochemical quenching of chlorophyll fluorescence. (m-r) Effect of range of concentrations of rifampicin upon the circadian oscillation of the apparent quantum yield of PSII (Y(II)). (s-x) Effect of range of concentrations of rifampicin upon the circadian oscillation of non-photochemical quenching of chlorophyll fluorescence. (e, f, k, l, q, r, w, x) Relative amplitude error, period and amplitude were estimated using FFT-NLLS with amplitude and baseline detrending. (e, k, q, w) Different letters next to data points indicate significant differences in period (upper case)) and RAE (lower case), with data plotted ± s.e.m. (f, l, r, x). Analyzed with one-way ANOVA, where p < 0.05 based on Tukey’s post-hoc test; n = 25 thalli. On time-series plots, white sections indicate subjective day and grey shading indicates subjective night.

Next, we tested the hypothesis that an inhibitor of chloroplast transcription can attenuate circadian rhythms of photosynthesis in *M. polymorpha.* In Arabidopsis, mutants of a circadian-regulated sigma submit of plastid encoded plastid RNA polymerase (PEP) known as SIG5 can alter the phase of circadian rhythms of delayed fluorescence (Noordally *et al.* 2013). Therefore, we reasoned that chloroplast transcription by PEP might contribute to circadian rhythms of photosynthesis in *M. polymorpha.* To test this, we supplemented the growth media with rifampicin, which is thought to inhibit chloroplast transcription by one form of PEP (Pfannschmidt and Link 1997, Surzycki 1969). We measured rhythms of Y(II) and NPQ, under free running light conditions, in the presence of several concentrations of rifampicin (Fig. 4m-x; Fig. S4e-h). Across two experiments, under these experimental conditions, there were no consistent differences caused by the range of rifampicin concentrations tested upon the circadian rhythm of the apparent quantum yield of PSII or NPQ (Fig. 4q, r, w, x; Fig. S4e, f, g, h).

## Discussion

Here, we present evidence for the circadian regulation of two proxies for photosynthetic activity in *M. polymorpha.* Our experiments cannot prove that circadian rhythms of photosynthesis occurred in early land plants, but the presence of these rhythms in both bryophytes and angiosperms suggests that they might be well-conserved. Given that reporters of photosynthetic activity are also circadian regulated in aquatic photosynthetic organisms such as cyanobacteria, uni- and multicellular algae (Cano-Ramirez *et al.* 2018, Hastings *et al.* 1961, Okada *et al.* 1978, Schneegurt *et al.* 1994, Schweiger *et al.* 1964, Sorek and Levy 2012, Sorek *et al.* 2013, Sweeney and Haxo 1961), our study suggests that the circadian regulation of photosynthesis was conserved from aquatic ancestors during the terrestrialization of plants. One interpretation is that circadian regulation of photosynthesis confers a selective advantage to all photosynthetic life. The nature of the advantage and degree of selection pressure for this might depend upon the ecological niche occupied by the organism (Hellweger *et al.* 2020), particularly because certain species (e.g. *Picea abies)* appear to lack these rhythms (Gyllenstrand *et al.* 2014).

Our experiments identified that under free running light conditions, the period of the rhythm of delayed fluorescence was not uniform between replicate thalli, with this variation leading to phase desynchrony (Fig. 1a-d). In comparison to *M. polymorpha,* the circadian rhythm of DF in Arabidopsis and other flowering plants is relatively stable, with a period of about 24 h and individual replicates relatively synchronized (Dataset S1) (Gould *et al.* 2009, Rees *et al.* 2021). The range of phases of Y(II) for replicate thalli, under free running conditions, was also rather broad because the phases were spread across a period of 8 h within the 24 h cycle (Fig. S6). Other reporters for circadian rhythms in *M. polymorpha* are also less robust than in flowering plants. For example, rhythms of some promoter-luciferase reporters damp quite rapidly and present a range of periods across replicates, with this difference in robustness evident when circadian rhythms in *M. polymorpha* and Arabidopsis are compared (Lagercrantz *et al.* 2021, Linde *et al.* 2017) (Dataset S1). This desynchronization of replicate thalli is reminiscent of the phase desynchronization that occurs between the rhythms of individual cells in the leaves of *Lemna gibba* under free running conditions, compared with the maintenance of phase synchrony in *L. gibba* under zeitgeber cycles (Muranaka and Oyama 2016). Each of these experiments occurred under different light conditions, with DF and PAM measurements involving fluctuations in the light conditions, so comparisons require a degree of caution because the stability of the *M. polymorpha* circadian system might vary considerably according to the light quality and quantity.

One potential explanation for this difference between circadian rhythms in *M. polymorpha* and flowering plants might relate to circadian oscillator structure (Linde *et al.* 2017). The circadian oscillator in *M. polymorpha* has been proposed to include a smaller number of components than the oscillator of Arabidopsis, with greater levels of oscillator complexity thought to confer robustness to perturbation (Shalit-Kaneh *et al.* 2018, Troein *et al.* 2009). Therefore, the less robust circadian oscillations of photosynthesis that we report might arise from lower levels of redundancy within a less complex circadian clock in *M. polymorpha* compared with Arabidopsis. An alternative explanation for this difference might relate to differences in the strength of coupling of circadian rhythms between cells and across tissues in Arabidopsis and *M. polymorpha.* Although there is weak intercellular coupling of circadian rhythms in Arabidopsis (Wenden *et al.* 2012), differences between the circadian period of individual cells means that the rhythms of cell populations become progressively more desynchronized over time in the absence of intercellular coupling, leading to an apparent damping of the rhythm (Gould *et al.* 2018, Muranaka and Oyama 2016, Paajanen *et al.* 2021). Therefore, rhythms might damp more rapidly in *M. polymorpha* if intercellular coupling of circadian rhythms is weaker compared with Arabidopsis, especially when combined with a potentially less robust circadian oscillator within each cell. The idea of relatively unstable circadian rhythms in the bryophytes is also supported by DF data from *Physcomitrium patens,* which is arrhythmic under constant light and has a noisy rhythm under low light conditions (Gyllenstrand *et al.* 2014). It will be interesting in future to determine whether there is simply phase desynchronization between individual replicate thalli of *M. polymorpha,* or patchy or complete intercellular desynchronization within individual thalli.

The thallus of *M. polymorpha* lacks stomatal pores for gas exchange and, instead, has air pores in its surface that are formed by a channel passing through a cylindrical structure of 16 cells (Jones and Dolan 2017, Shimamura 2016). Unlike the stomata of flowering plants, the air pores of *M. polymorpha* are thought to be static structures that do not adjust in response to environmental cues. Therefore, circadian rhythms of the reporters of photosynthesis described here might be due entirely to the circadian regulation of the biochemical reactions of photosynthesis, rather than changes in the rate of CO_2_ and O_2_ exchange with the atmosphere. Nevertheless, we cannot rule out alterations in conductivity to gas exchange through diel or circadian changes in cell turgor or potential “motor cells” within liverwort air pores (Walker and Pennington 1939). Such changes might influence the measures of photosynthesis that we used, because the presence of the pores and air spaces within the thallus appears to reduce its resistance to CO2 diffusion (Green and Snelgar 1982).

The reduction in robustness of rhythms of Y(II) caused by lincomycin could suggest that chloroplast translation is necessary for circadian rhythms of this reporter of photosynthesis. Lincomycin inhibits the synthesis of PSII D1, PSII D2 and RbcS proteins, and the expression of photosynthesis-associated nuclear encoded genes (PhAnGs) in angiosperms (Bachmann *et al.* 2004, Karpinska *et al.* 2017, Mulo *et al.* 2003). Perhaps inhibiting the expression of these proteins prevents the replacement of PSII components, with ensuing impacts upon circadian cycles of light harvesting. The absence of any effect of rifampicin on circadian rhythms of Y(II) or NPQ, contrasting the effect of lincomycin, could suggest that rhythms of these parameters involve genes transcribed by nuclear encoded plastid RNA polymerase (NEP) rather than rifampicin-sensitive PEP, there is compensation by NEP for reduced PEP activity, or specific roles for the rifampicin sensitive- and insensitive forms of PEP (Pfannschmidt and Link 1994). Whilst this suggests that protein synthesis in chloroplasts is necessary for the circadian regulation of photosynthesis, it does not identify that this is a rate-limiting regulatory step that underlies the clock control of the process. It also supports the idea that *M. polymorpha* represents an informative model for investigation of mechanisms underlying the circadian regulation of physiology, particularly considering the range of gene-editing tools available.

Whilst the apparent reduction of robustness of circadian rhythms in *M. polymorpha* might increase the susceptibility of rhythms in *M. polymorpha* to environmental perturbation (Linde *et al.* 2017), it is alternatively possible that a circadian oscillator with greater variability might confer selective advantages. For example, plasticity of circadian regulation of the daily photosynthetic cycle of CAM plants, and of carbohydrate metabolism in Arabidopsis, are both thought to allow better alignment between metabolic processes and daily cycles of energy availability (Borland *et al.* 2011, Dodd *et al.* 2002, Graf *et al.* 2010, Webb *et al.* 2019). In comparison, very little is known about diel cycles of metabolism in *M. polymorpha,* or how circadian regulation might confer fitness to *M. polymorpha.* To provide a better context for understanding these findings, it might be valuable in future to investigate the regulation and adaptive functions of the circadian system in *M. polymorpha,* or in naturally occurring populations of *M. polymorpha.*

## Materials and Methods

### Plant material and growth conditions

Tissue of *Marchantia polymorpha* subsp. *ruderalis* (Tak-1 background) was cultivated under sterile conditions, using half-strength Gamborg B5 media containing 0.8% (w/v) agar at pH 5.8 (Solly et al. 2017). Cultivation occurred in Panasonic MLR-352 growth chambers, under cycles of 12 h light / 12 h darkness, at 19 °C, under 100 μmol photons m^-2^ s^-1^ of white light (light spectrum in Fig. S5a). For inhibitor experiments, lincomycin (Alfa Aesar) or rifampicin (Melford Laboratories Ltd) were added for the PAM chlorophyll fluorescence experiments at concentrations of 25, 50 and 100 μg/ml, with a dimethylsulfoxide (DMSO) vehicle control where necessary. In *M. polymorpha,* rifampicin is known to inhibit plastid rRNA transcription (Loiseax *et al.* 1975), but does not inhibit photosynthesis (Mache and Loiseax 1972). The effect of lincomycin upon *M. polymorpha* is less well characterized, although it is known to bind to 50S ribosomes (Chang and Weisblum 1967), which are present in chloroplasts of *M. polymorpha* (Ohyama *et al.* 1986).

For DF experiments, gemmae (Shimamura 2016) were positioned on nutrient agar and cultivated for 14 days prior to the start of the experiment, and entrained under 12 h light / 12 h dark cycles for this entire period. For PAM fluorescence experiments, pieces of 21-day old thallus cultivated under 12 h light / 12 h dark cycles were cut into square sections under sterile conditions and allowed to regenerate for 6 days, during which they were entrained under 12 h light / 12 h dark cycles. Preparation of the thallus as square shapes prevented hyponasty during these experiments. For both DF and PAM experiments, plants were entrained under conditions described above prior to imaging.

### PAM chlorophyll fluorescence measurement

Circadian time-series of chlorophyll fluorescence measurements were obtained using an IMAGING-PAM M series chlorophyll fluorescence imaging system (Walz GmbH, Effeltrich, Germany), using a protocol described previously (Cano-Ramirez *et al.* 2018). Briefly, rapid light curves (RLCs) were obtained at 2 hour intervals, with the plants maintained under constant blue light at other times during the timecourse. Each RLC was acquired over a period of 6 minutes, which was preceded by a 20 minute period of dark adaptation. For the acquisition of each RLC, a saturating light pulse (845 μmol m^-2^ s^-1^) was applied, and followed by an increase in actinic light across 16 steps, at 20 second intervals, to a maximum of 789 μmol m^-2^ s^-1^. After acquiring the RLC, samples were held under continuous blue light (15 μmol m^-2^ s^-1^; light spectrum in Fig. S5b) until the next measurement cycle. PAM chlorophyll fluorescence parameters were measured at 108 μmol m^-2^ s^-1^ actinic light on the RLC, which was similar to the growth and entrainment conditions (100 μmol m^-2^ s^-1^).

### Measurement of delayed fluorescence

Delayed fluorescence analysis was performed as described previously (Gould *et al.* 2009, Rees *et al.* 2019). This involved image capture with Retiga LUMO cameras (QImaging, Canada), fitted with Xenon 0.95/25 mm lenses (Scheneider-Kreuznach, Germany). Plants were illuminated with mixed red/blue LED arrays (approx. 60 μmol m^-2^ s^-1^; light spectrum in Fig. S5c) under the control of the μManager Software (v1.4.19, Open Imaging) and an Arduino Uno microcontroller board. The camera and LEDs were mounted in growth chambers maintained at 19 °C (Sanyo MIR-553). The camera settings for all experiments were Binning = 4, Gain = 1, and Readout-Rate=0.650195 MHz, 16 bits. Three control scripts were used for DF measurement in *Marchantia polymorpha.* The free running light condition script was a regime of 59 minutes of light followed by one minute of image acquisition, the free running dark condition script was 54 minutes of darkness followed by five minutes of light and one minute of image acquisition, and the zeitgeber script was an alternating combination of FRL and FRD scripts that swapped every half T-cycle. Exposure of plants to 5 minutes of light before image acquisition under both free running dark conditions and the dark period of zeitgeber cycles was necessary to obtain the DF signal, and might be perceived by plants as a low fluence response equivalent to an average photosynthetically active radiation (PAR) of approximately 3 μmol m^-2^ s^-1^ over each hour. All experiments started at entrained dawn, and finished after seven days of imaging. T-cycle experiments employed the zeitgeber cycle script, but using daily periods of 20 h (10L/10D), 22 h (11L/11D) and 28 h (14L/14D), respectively. For analysis, images were imported into FIJI (version 1.53c) and regions of interest were selected and extracted using the multi-measure plugin to measure integrated density of each region. Raw data before filtration are provided in Dataset S2. The DF data were baseline and amplitude (BAMP) de-trended using Biodare2, and a moving average was used to smooth traces from individual thalli.

### Data analysis

The DF data were smoothed using a 5 h moving average (Gould *et al.* 2009) and baseline and amplitude (BAMP) detrended using Biodare2 (Zielinski *et al.* 2014). These data are provided Fig. 1b, d, f and Fig. S1, S2 and S3 to exemplify variation in the DF rhythms across all thalli. Data detrending was necessary because rapid thallus growth during the experiment produced a continuously increasing raw signal. The data were subsequently analyzed using Metacycle (Wu *et al.* 2016) to identify rhythmic thalli (*q* < 0.001; period range 18 – 34 h). Data that passed this filtration step were used for calculation of rhythmic features and presented in Fig. 1a, c, e, g-k and Table 1.

Rhythmic features of the data were quantified using the Fast Fourier Transform-Non-linear least squares method (FFT-NLLS) algorithm within Biodare2 (Zielinski *et al.* 2014), and also with Metacycle (Wu *et al.* 2016). Statistical analysis on derived data was conducted using IBM SPSS Statistics v24 and SigmaPlot v14 for PAM chlorophyll fluorescence and DF data. Graphs were prepared using R, with a combination of the ggplot2, scales, rhesape2, cowplot, png, grid, RColorBrewer, wesanderson, colorspace, ggpubr, ggrepel, and patchwork packages. Heat maps were created using the R package Pheatmap. A 5 h moving average was applied to DF and PAM fluorescence data before analysis (Gould *et al.* 2009).

For calculation of the change in mean RAE and synchronisation index over time (Fig. 1j, k), data were smoothed using a moving average, and filtered using Metacycle to identify rhythmic thalli *(q* <0.001; period range 18 – 34 h). RAE estimates and phases were recalculated using FFT-NLLS across a 48h sliding window (e.g. 24-72 h, 48-96 h, and so on) Only rhythmic thalli that did not produce error messages within Biodare were included in the means for each time window (FRL: N= 35-52; FRD: N=47-62; ZTC: N=95-98). The synchronisation index is Kuramoto’s order parameter, *r* (Kuramoto 1984, Nykamp accessed 2022), which represents the synchrony of phases in any given time window, where *r* = 1 indicates absolute phase synchrony. Our code for the calculation of the synchronisation index and mean phases for each time window is available on GitHub (https://github.com/AHallLab/KuramotoParameters).

## Acknowledgements

We thank our funders: UKRI-BBSRC Institute Strategic Programme (ISP) Genes in the Environment (BB/P013511/1), Core Strategic Programme Grant 901 (Genomes to Food Security BB/CSP1720/1) and its workpackage 902 BBS/E/T/000PR9819 (Regulatory Interactions and Complex Phenotypes), Designing Future Wheat ISP (BB/P016855/1; 904 BBS/E/T/000PR9783 (WP4 Data Access and Analysis), and Conacyt (Mexico) (CVU: 510730, Scholarship: 472284). We thank Jill Harrison (Bristol), Ryuichi Nishihama (Kyoto) and Takayuki Kohchi (Kyoto) for advice about experimentation with *M. polymorpha,* Martha Merrow (LMU), Zheng Eelderink-Chen (LMU) and Jasper Bosman (Hanze Univ.) for advice about data analysis and interpretation, and Maria Paola Puggioni, Aurélie Crepin and colleagues (Umeå University) for biorxiv feedback.

---

**Figure S1.** Example of raw delayed fluorescence data for 64 individual replicate thalli of *M. polymorpha*, under free running light conditions. Clear areas and shaded areas on the plots indicate subjective day and night, respectively.

**Figure S2.** Example of raw delayed fluorescence data for 64 individual replicate thalli of *M. polymorpha*, under free running dark conditions. Clear areas and shaded areas on the plots indicate subjective day and night, respectively.

**Figure S3.** Example of raw delayed fluorescence data for 64 individual replicate thalli of *M. polymorpha*, under zeitgeber cycles. Clear areas and shaded areas on the plots indicate day and night, respectively.

**Figure S4.** Amplitude of circadian rhythm of chlorophyll fluorescence parameters is reduced by the inhibitor of chloroplast translation lincomycin. This is a second direct repeat of the experiment in Fig. 4. (a-d) Effect of range of concentrations of lincomycin upon the circadian oscillation of (a, b) the apparent quantum yield of PSII (Y(II)) and (c, d) non-photochemical quenching of chlorophyll fluorescence. (e-h) Effect of range of concentrations of rifampicin upon the circadian oscillation of (e, f) Y(II) and (g, h) non-photochemical quenching of chlorophyll fluorescence. (a, c, e, g) Different letters next to data points indicate significant differences in period (upper case) and RAE (lower case), with data plotted ± s.e.m. (b, d, f, h) Different letters indicate significantly different mean phases. Analyzed with one-way ANOVA, where p < 0.05 based on Tukey’s post-hoc test; n = 25 thalli.

**Figure S5.** Spectra of light conditions used for experiments. Photon flux density spectrum within (a) growth chambers used for cultivation and entrainment of M. polymorpha, (b) imaging PAM system actinic light conditions and (c) delayed fluorescence imaging LED array.

**Figure S6.** Phase distribution for Y(II) in *M. polymorpha* under free running conditions measured using PAM fluorescence. Phases of replicate thalli relative to dawn (0), calculated using fast Fourier transform-non linear least squares method (FFT-NLLS). Radial scale shows relative amplitude error (RAE) of each thallus, with RAE=0 at the exterior and RAE=1 at the centre of the circles, respectively.

**Dataset S1.** Comparison of synchrony of circadian rhythms of delayed fluorescence in variety of species and experimental conditions. This uses using the order parameter within the Kuramoto phase oscillator model to provide a synchronization index. Data derive from this paper, (Rees *et al.* 2019, Rees *et al.* 2021).

## Notes

### Competing Interest Statement

The authors have declared no competing interest.

### Summary of Updates

Small revisions in response to reviewer and editorial comments, and addition of further supplemental figure.

