## Supplementary figures and images for "Circadian and diel regulation of photosynthesis in the bryophyte *Marchantia polymorpha*"

### Figure S1

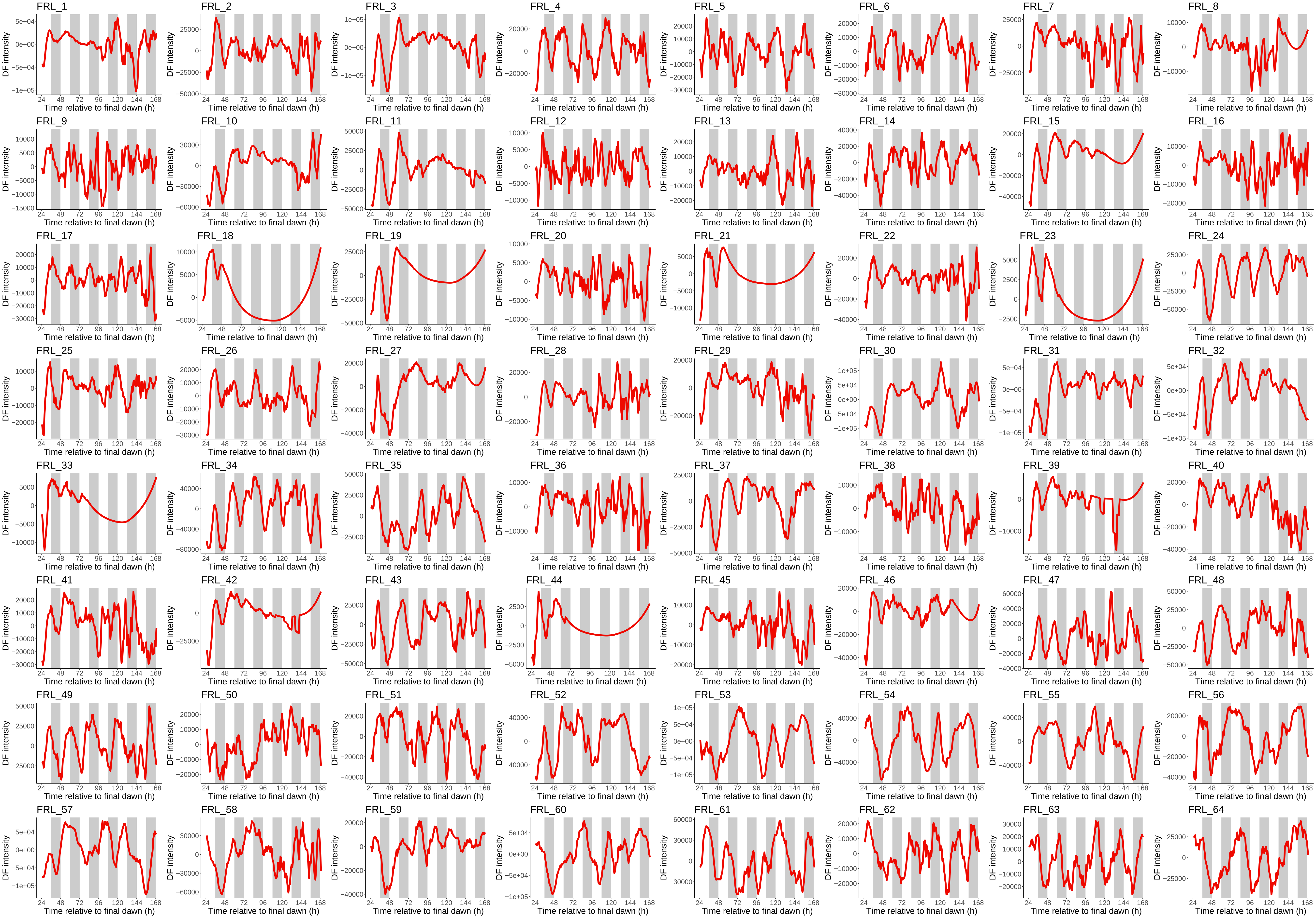

### Figure S2

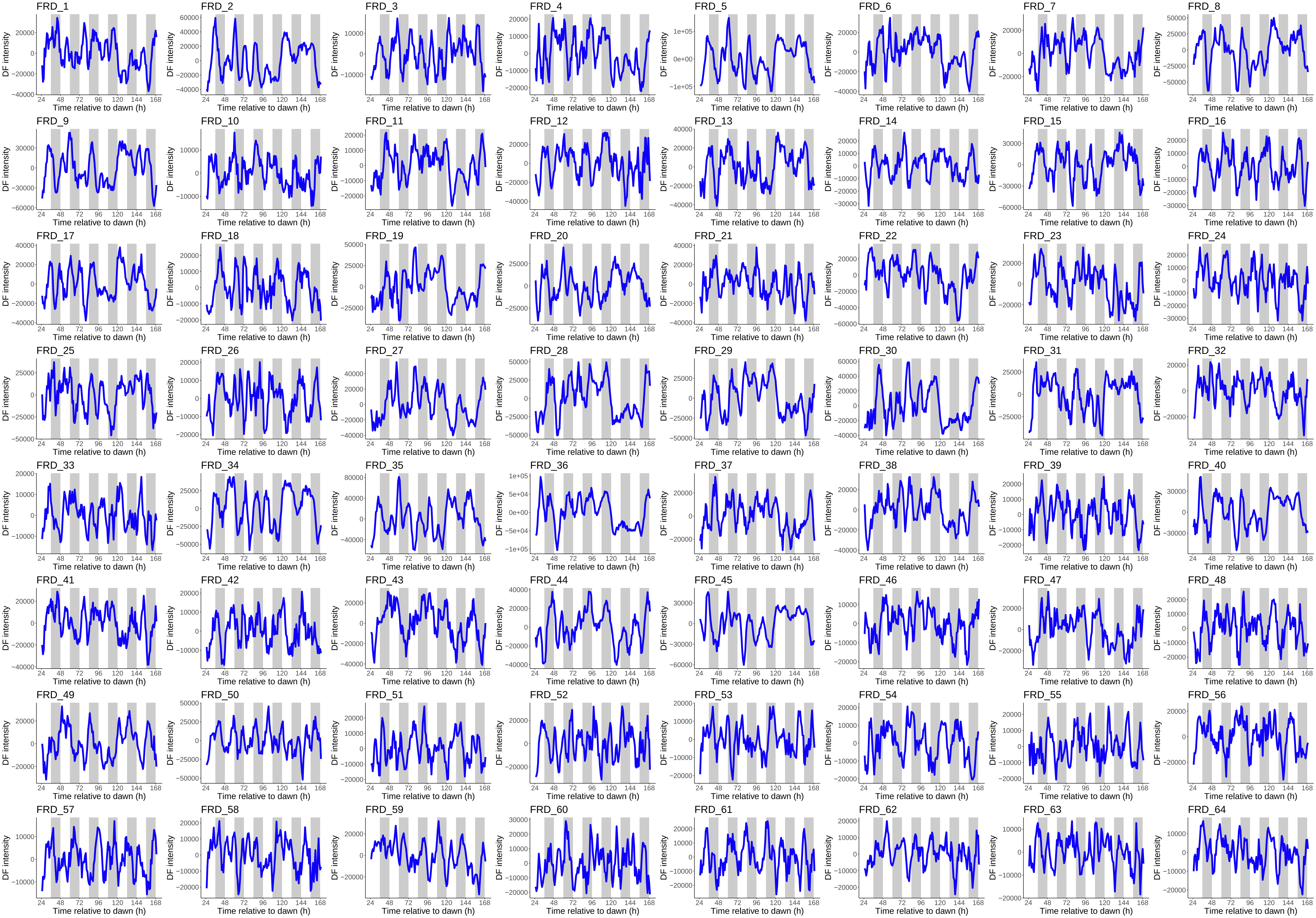

### Figure S3

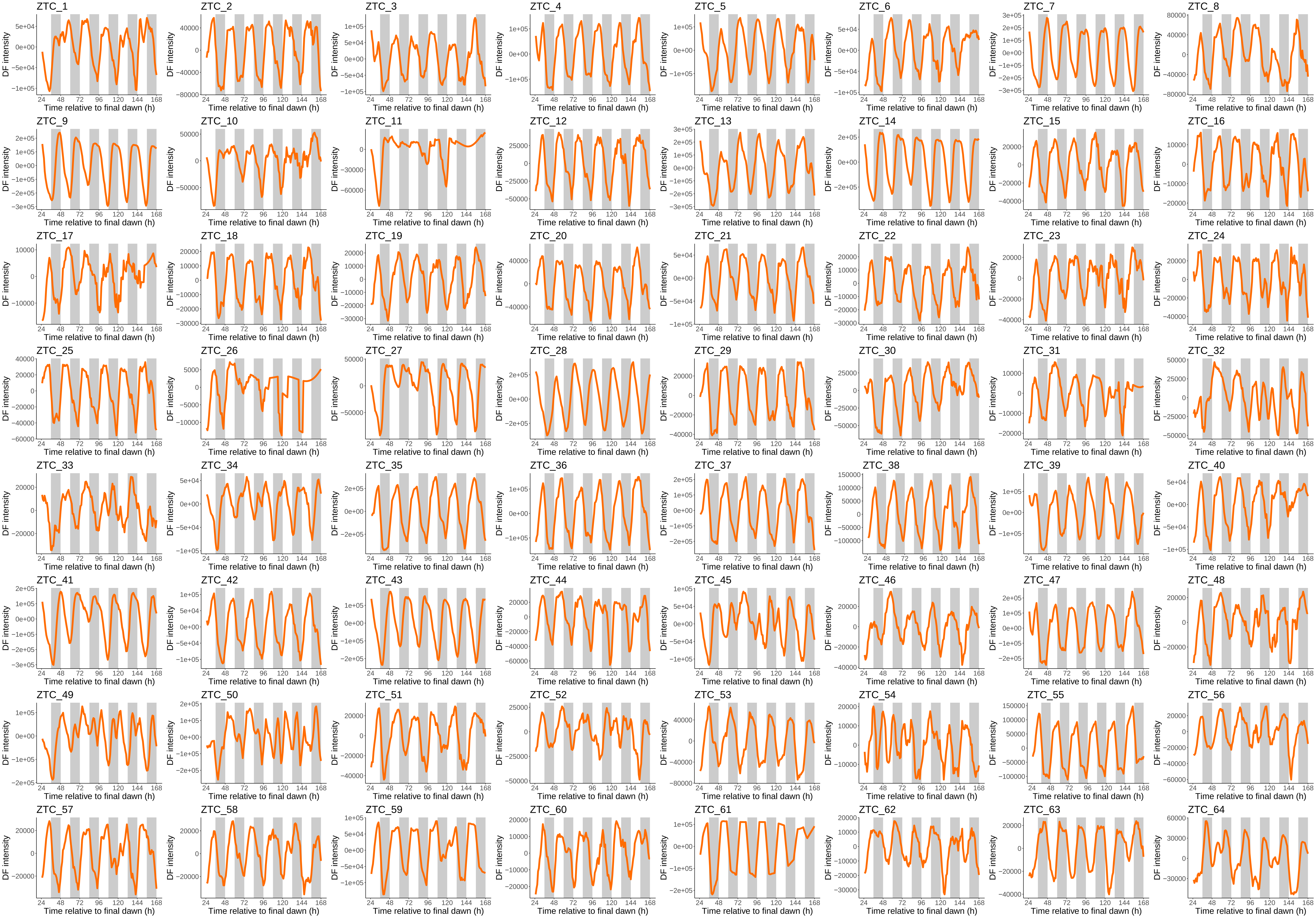

### Figure S4

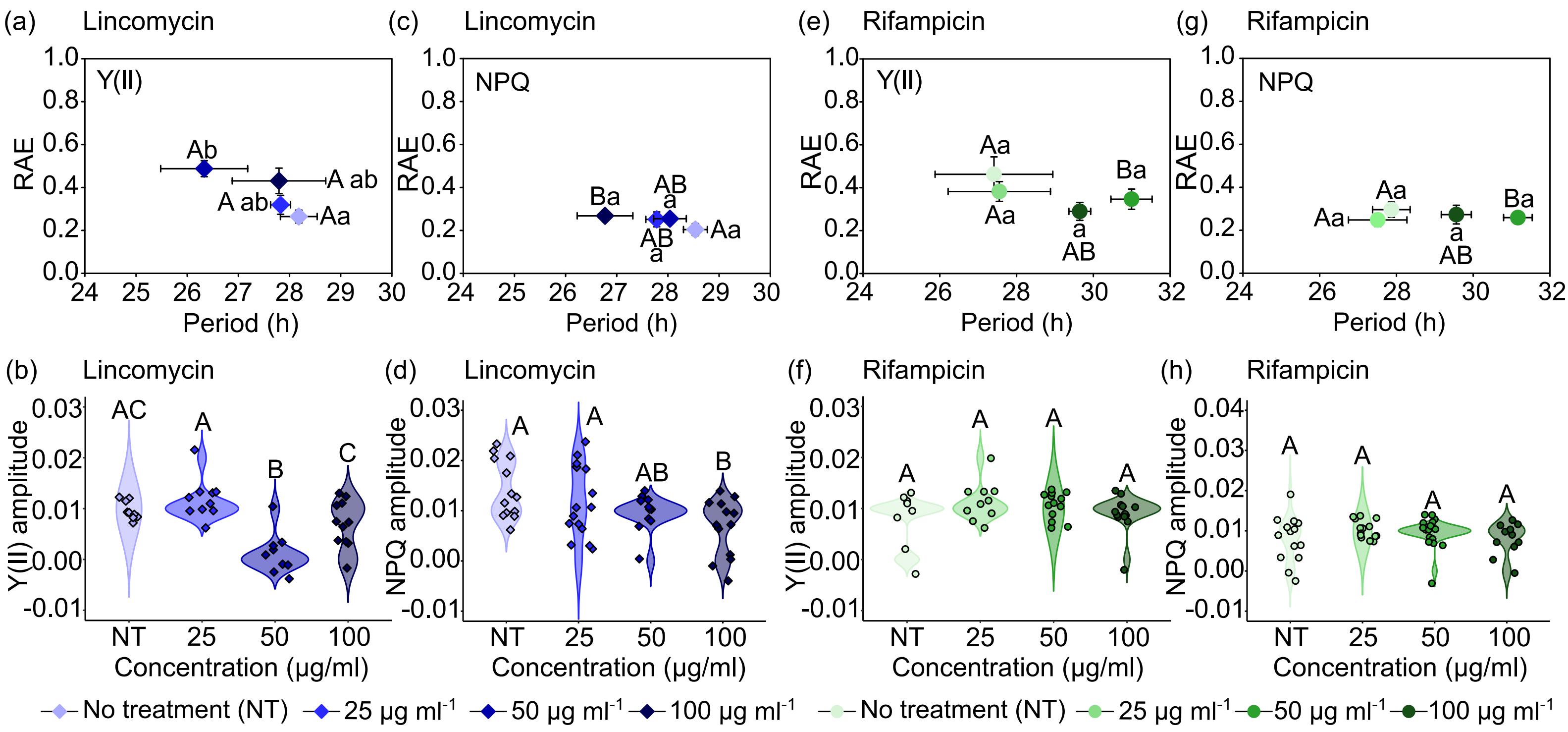

### Figure S5

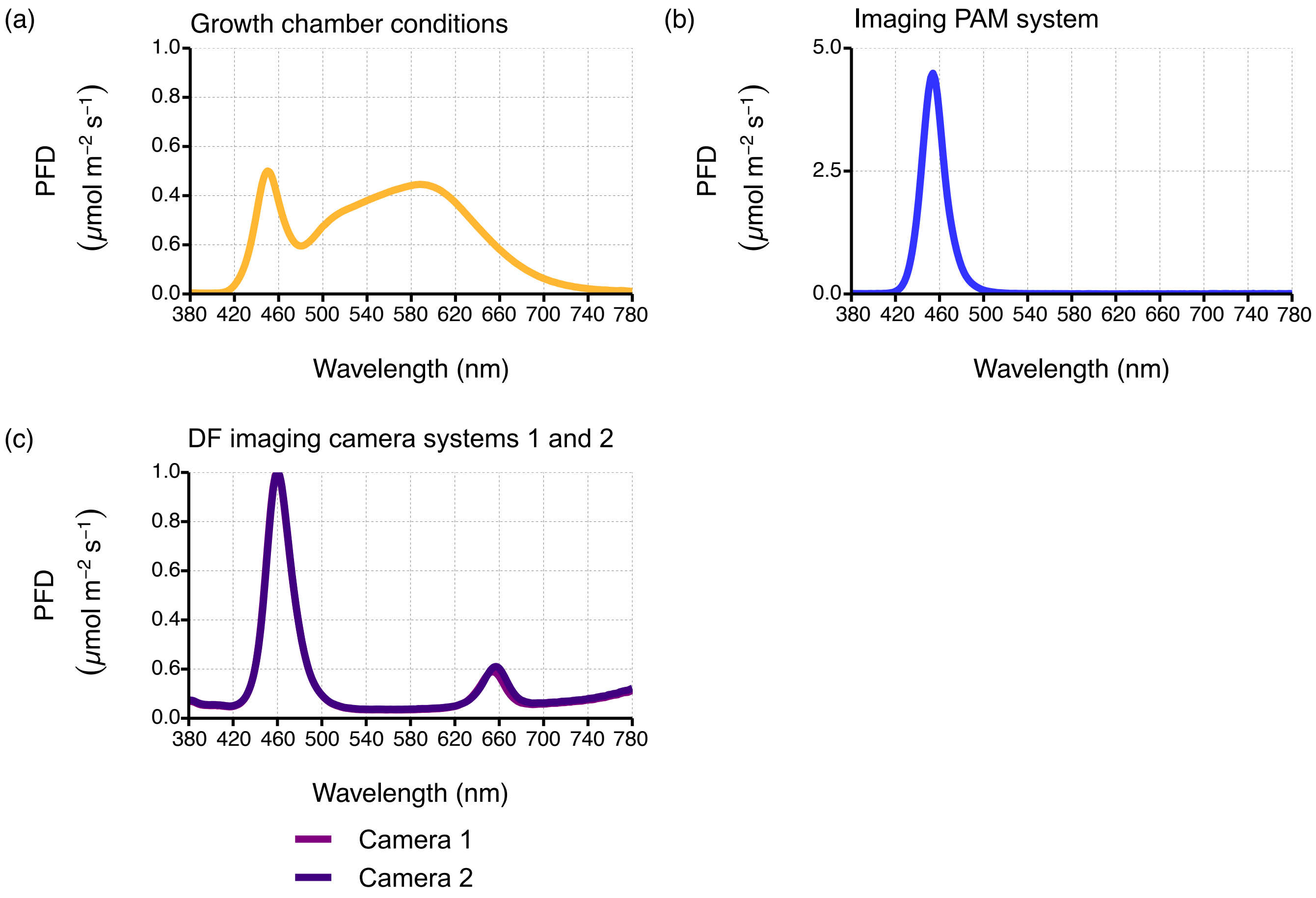

### Figure S6

Y(II)

Free running conditions

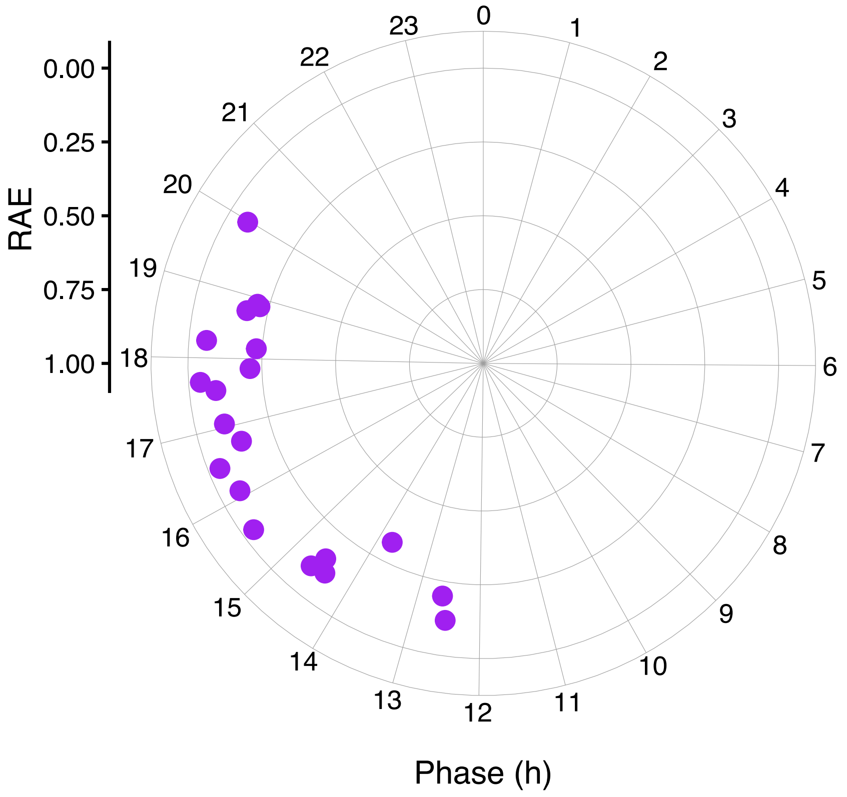
